# Interpreting Deep Neural Networks Beyond Attribution Methods: Quantifying Global Importance of Genomic Features

**DOI:** 10.1101/2020.02.19.956896

**Authors:** Peter K. Koo, Matt Ploenzke

## Abstract

Despite deep neural networks (DNNs) having found great success at improving performance on various prediction tasks in computational genomics, it remains difficult to understand why they make any given prediction. In genomics, the main approaches to interpret a high-performing DNN are to visualize learned representations via weight visualizations and attribution methods. While these methods can be informative, each has strong limitations. For instance, attribution methods only uncover the independent contribution of single nucleotide variants in a given sequence. Here we discuss and argue for global importance analysis which can quantify population-level importance of putative features and their interactions learned by a DNN. We highlight recent work that has benefited from this interpretability approach and then discuss connections between global importance analysis and causality.

## Overview

Deep neural networks (DNNs) have demonstrated improved performance in many prediction tasks in computational biology (Zhou & Troyanskaya, 2015; Alipanahi et al., 2015; Zeng et al., 2016; Eraslan et al., 2019; Hiranuma et al., 2017; Angermueller et al., 2017; Kelley et al., 2016). Despite their promise, the main drawback of DNNs is the difficulty in understanding why they make any given prediction. Treated as a black box, it is challenging to decipher whether improved predictions result from learning novel biological features not captured by previous methods or by gaining an advantage through discriminating correlated features that are indirectly related, such as technical biases of an experiment. Models that exploit the latter may not necessarily generalize well, especially across datasets generated by different protocols, laboratories, or sequencing technologies.

Currently, the main approach to interpret a convolutional neural network (CNN) is to visualize learned representations in the input space. In genomics, such methods include visualizing the convolutional filters (Alipanahi et al., 2015; Kelley et al., 2016; Quang & Xie, 2016; Angermueller et al., 2016; Cuperus et al., 2017; Chen et al., 2018; Ben-Bassat et al., 2018; Wang et al., 2018), attribution methods (Alipanahi et al., 2015; Zhou & Troyanskaya, 2015; Kelley et al., 2016; Shrikumar et al., 2017; Lundberg & Lee, 2017; Ghanbari & Ohler, 2019), and more recently *in silico* experiments (Koo et al., 2018; Avsec et al., 2019). These approaches can be grouped into *local* and *global* interpretability methods. Local interpretability methods provide sample-level feature importance, that is for individual sequences, while global interpretability methods describe population-level feature importance. Here, we give a brief overview of local and global interpretability methods and then argue for the latter. We highlight two applications where global interpretability of a high-performing DNN has provided a more in-depth understanding of the underlying biology.

## Local interpretability

In genomics, attribution methods – such as *in silico* mutagenesis (Alipanahi et al., 2015; Zhou & Troyanskaya, 2015; Kelley et al., 2016), gradients to inputs, *i.e*. saliency maps (Simonyan et al., 2013), integrated gradients (Sundararajan et al., 2017), and Deeplift (Shrikumar et al., 2017) – provide a nucleotide-resolution map consisting of an importance score for each nucleotide variant at each position for a given sequence. There are several other interpretability methods that have not been thoroughly explored in regulatory genomics applications, including deconvolution (Zeiler & Fergus, 2014), GRAD-CAM (Selvaraju et al., 2017), SHAP (Lundberg & Lee, 2017), and LIME (Ribeiro et al., 2016), among others not cited here. The main benefit of attribution methods is that they provide importance scores related to decisions, considering the entire DNN.

In practice, many applications have utilized attribution methods to validate that their model has learned meaningful biology. For example, gradients (from predictions to the inputs) have been employed to reveal known transcription factor (TF) binding sites when trained to predict read profiles from high-throughput sequencing datasets (Kelley et al., 2018). Integrated gradients were used to uncover motifs for RNA-protein interactions (Ghanbari & Ohler, 2019). Recently, DeepLift was used to uncover known and novel TF binding sites, including their syntax with respect to other binding sites (Avsec et al., 2019). *In silico* mutagenesis - the gold standard for local interpretability in genomics - has been shown to uncover known motifs related to TF binding and chromatin accessibility (Alipanahi et al., 2015; Zhou & Troyanskaya, 2015; Kelley et al., 2016). Collectively, these approaches have been useful to validate DNN predictions for known disease-associated variants, albeit on an anecdotal basis. More recently, local interpretability has helped to understand the role of noncoding mutations in autism spectrum disorder and to prioritize high-impact mutations for further study (Zhou et al., 2019).

## Global interpretability

Although attribution methods can be informative, their main drawback is that it can only be applied *locally* to individual sequences. However, any putative patterns that are identified in a given sequence may be influenced by other factors, such as the 3D structure of the sequence or interactions with other proteins. It remains difficult to disentangle whether attribution scores are noisy due to an artifact of the attribution method itself or a consequence of poor representations learned by the DNN. To convert sample-level representations captured locally in attribution maps into global representations at the population-level, TF-MoDISco splits attribution maps into smaller segments about learned patterns called seqlets and clusters these seqlets to find averaged representations, which reduces noise from any individual seqlet (Shrikumar et al., 2018). Alternatively, global feature representations can be identified by visualizing first layer convolutional filters. This can be accomplished by directly plotting their weights or via activation-based sequence alignments, which are converted to a position frequency matrix. Recent advances have made it possible to intentionally design CNNs to learn more human-interpretable patterns in convolutional filters. This includes design principles based on spatial information flow through the network and employing highly divergent activation functions such as the exponential function (Koo & Eddy, 2019; Koo & Ploenzke, 2019). In parallel, advances have been developed to make direct weight visualization more interpretable (Ploenzke & Irizarry, 2018).

Visualizing convolutional filters has the benefit of revealing *global* sequence features. However, information about interactions between first layer features is captured in the deeper layers. For CNNs that employ pooling, deeper layers cannot be visualized by the standard activation-based alignments because spatial information of filter activations is lost after each pooling operation. Another drawback is the lack of correspondence between first layer features to decision making (the output layer). TF-MoDISco uses attribution maps, so it should, in principle, provide class-specific representations. Nevertheless, it is still unable to quantify the importance or effect size of the class-specific features. Recently, it was shown that networks that make state-of-the-art predictions can still yield unreliable attribution scores (Tsipras et al., 2018; Alvarez-Melis & Jaakkola, 2018; Koo et al., 2019). Thus, there are no guarantees that attribution maps in genomics will provide biologically meaningful representations just because the DNN makes accurate classifications.

### Global importance analysis: a quantitative interpretability approach

Heretofore, interpretability methods have been primarily used in genomics to show that a DNN has learned representations that match previously known motifs, serving as validation for its performance. While attribution methods provide a quantitative importance score for individual nucleotide variants, they do not provide the statistical importance of any putative features, like motifs. To quantitatively uncover the *global* importance of a motif, we would ideally average the model predictions over a corpus of sequences which contain the motif under investigation, while also randomizing the other positions so that background noise and extraneous confounding signals may be allayed. Mathematically, this is expressed as:

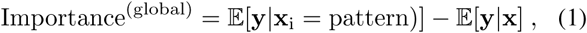

where 𝔼 is an expectation, **y** are the network predictions for input sequences **x**, and **x**_*i*_ represents the input sequences with the studied pattern embedded starting at the *i*th position. Equation 1 quantifies the effect size of just the embedded feature at a specific position by marginalizing out the contributions of the other positions.

Important to this approach is the randomization of all other positions. Since the necessary sequences to calculate Eq. 1 may not exist in a given dataset, one solution is to generate synthetic sequences. Such a procedure requires selection of an appropriate sequence model to minimize any distribution shift between the synthetic sequences and the experimental data. One approach can be to sample sequences from a profile based on a site-independent sequence model. Here, one would expect that the profile model captures all position-dependent biases that are present across the entire experimental dataset but not any position-independent patterns, like motifs. Alternative null models include random shuffling and dinucleotide shuffling of the real sequences in the dataset. If there exists high-order dependencies in the observed sequences, such as RNA secondary structure or motif interactions, or if background features do not have a strict positional dependence, a distribution shift may arise between the null model and data distribution, which can lead to misleading results. In genomics, prior knowledge can help design a suitable null sequence model. Another strategy can be to “knocked-in” candidate motifs into real genomic sequences to measure the effect size that the motif has on model predictions. Suitable genomic sequences may come from the negative class in binary classification tasks.

Global importance analysis can be employed to uncover higher-order interactions by embedding two (or more) candidate motifs in null model sequences and varying their spacing. Occluding regions or patches in the data is another powerful method to discover important features in images (Zeiler & Fergus, 2014). In genomics, this is analogous to an *in silico* CRISPR experiment.

### Beyond validation - discovering new biology

To the authors knowledge, (Koo et al., 2018) was the first demonstration of interpreting a DNN in genomics using global importance analysis. They employed global importance analysis to show that their DNN, called ResidualBind, trained to infer sequence-structure preferences of RNA binding proteins (RBPs), learned not only the underlying sequence motifs, but also based predictions on the number of motifs, their spacing, and secondary structure context. At the time, other DNNs had been applied to the same RNAcompete dataset (Ray et al., 2013), including Deepbind (Alipanahi et al., 2015), DeeperBind (Hassanzadeh & Wang, 2016), DLPRB (Ben-Bassat et al., 2018), and cDeepbind (Gandhi et al., 2018). Each method benchmarked their performance on held out test data from RNAcompete and also on *in vitro*-to-*in vivo* generalization tasks. For interpretability, Deepbind and DLPRB demonstrated that a few first layer convolutional filters learn representations that represent known RBP motifs. Deepbind and cDeepbind performed *in silico* mutagenesis anecdotally on a few sequences to show that their models have learned representations that resemble known RBP motifs.

In contrast, to interpret ResidualBind, (Koo et al., 2018) initially employed first-order *in silico* mutagenesis to show that canonical motifs are indeed learned. But this in itself does not explain ResidualBind’ s improved performance because previous methods also converged on similar motif representations. By performing *in silico* mutagenesis on a ResidualBind model trained on the RNAcompete dataset for RBFOX1, which has an experimentally validated motif ‘UGCAUG’, they were able to generate hypotheses that ResidualBind is learning to count the number of motifs, consider spacing between the motifs and their positions along the RNA probes. Using global importance analysis, they carried out *in silico* experiments to test each hypothesis. For instance, they systematically varied the number of RBFOX1 motifs in synthetic sequences to verify that ResidualBind integrates the presence of multiple binding sites in a given sequence with an additive model. They also varied the spacing between two RBFOX1 motifs in synthetic sequences to show that ResidualBind’ s predictions are consistent with a biophysical intuition of steric hindrance. They also interpreted a ResidualBind model trained on an RNA-compete dataset for VTS1, which has a sequence preference ‘GCUGG’ in the context of a hairpin loop. Using *in silico* mutagenesis, they found that the VTS1 motif was important for the network, but there were many other nucleotides that also exhibited high importance. These noisy positions were presumably features related to RNA secondary structure. To test this, they performed global importance analysis by designing synthetic sequences embedded with the VTS1 motif within the loop of a hairpin structure and the stem. Indeed the VTS1 motif had a statistically significant effect size in the hairpin loop. As a control, they embedded the VTS1 motif at similar positions in random sequences. These experiments support that ResidualBind has learned both positive and negative contributions of RNA structure context directly from the sequence despite never explicitly being trained to do so. Further, global importance analysis revealed that ResidualBind has learned a significant 3’ GC-bias for a subset of RBPs in the RNAcompete dataset.

Another demonstration of global importance analysis was in a recent study by (Avsec et al., 2019). They trained their DNN, called BPNet, to predict ChIP-nexus binding profiles for various transcription factors. To interpret BPNet, they first employed Deeplift, a local interpretability method, to quantify the contribution of each base pair in an input sequence. To summarize recurring patterns, they employed TF-MoDISco, a global interpretability method, to cluster Deeplift scores into motif representations called contribution weight matrices. They found 51 motifs, but focused on a subset of 11 TF motifs for further analysis, including the Nanog, Oct4, Sox2, and Klf4 motifs. They then performed global importance analysis to study properties of the learned motifs. Specifically, they designed *in silico* experiments in which they embed two TF motifs in synthetic sequences and systematically vary their separation. They found the Nanog motif was strongly enhanced by the presence of another Nanog motif nearby. Similar findings were noted for the Sox2 motif. Interestingly, they found directional dependencies in the enhancement of Nanog and Sox2 binding. They also performed occlusion experiments by removing motifs from real sequences and replacing them with random sequences. They found that Nanog motif instances exhibit a 10.5 basepair periodicity which corresponds to the helix property of DNA.

Together, these examples demonstrate the potential for interpreting high-performing models beyond local interpretability. Follow up global importance analysis can highlight patterns that are shared across the dataset, elucidating more informative representations that the model has learned, the specific function it has fit, and ultimately deeper insight into the underlying biology.

### Connection to causality

Recently, attribution methods for deep neural networks have been recast in a causal inference framework following the *do(.)* calculus (Chattopadhyay et al., 2019; Pearl, 2012). In this context, current attribution methods that are conditioned on a single data example are identifiable as a special instance of an individual causal effect (ICE). For a given data sample, 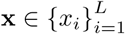 – where *x*_*i*_ is the *i*th feature and *L* is the number of input features – the ICE of setting the *i*th feature to a value *α* is estimated by: 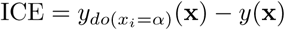, where 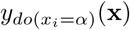 denotes the output *y* of a DNN when setting *x*_*i*_ to a value *α* and *y*(x) represents the DNN output for the unperturbed data sample. Setting a specific input feature to a value *α* is called an intervention and is represented with the *do* operation. Thus ICE estimates the effect size of an intervention to the *i*th feature for a given data sample. Employing an intervention with a small perturbation to a nucleotide variant is proportional to calculating the partial derivative with respect to the input, while intervening at the position level is similar to *in silico* mutagenesis. Systematically calculating ICE separately for each input feature generates an attribution map for a single sequence.

The causal effect of features identified by ICE are *local* to an individual data sample and hence may not necessarily generalize to the population level due to unaccounted feature interactions (endogenous confounders). To address this limitation, the average causal effect (ACE) calculates a feature’ s causal effect globally (Pearl, 2009), according to:

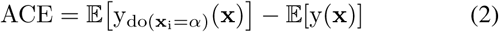

where 𝔼[·] is an expectation over the data distribution. For high-dimensional continuous random variables, such as images, ACE requires approximations to make the expectation tractable (Chattopadhyay et al., 2019).

In genomics, the causal effect of a specific sequence pattern with respect to a given molecular phenotype, such as protein binding, can be estimated by experimentally measuring the phenotypic outcome of sequences designed to contain a fixed, known pattern (intervention) and randomizing the other positions within the sequences as well as the intervention assignment. This process ensures ignorability of treatment assignment and a common support between treated and untreated, allowing for valid statistical inference of the causal effect. Equation 2 can thus be calculated directly with experimental measurements. In practice, this approach can be time consuming and costly due to the large number of sequences and experiments required to calculate Eq. 2. Alternatively, a well-trained neural network may be used as a surrogate for these “causal” experiments, generating predictions in lieu of experimental measurements for the sequences necessary to estimate Eq. 2. Indeed this is precisely what global importance analysis is doing.

Global importance analysis does not make any claims that it learns causal structure in the data. It is a tool that quantifies the effect size of patterns that are causally linked to model predictions. Hence, it should only be treated as a model interpretability tool. Although DNNs are only capable of learning associations, it may be possible that some of the associations play a causal role in terms of the data generating process. While model interpretability can suggest biological insights and help researchers to develop hypotheses, the patterns they learn are not proof of biological mechanisms. Any new insights made by interpreting a DNN should be followed up with wet lab experiments for validation.

## Conclusion

Global importance analysis is a powerful interpretability method that treats a high-performing DNN as an *in silico* laboratory, where *in silico* experiments can be carried out relatively cheaply. By leveraging randomized trials in the experimental design, we can quantitatively measure the effect size of putative features and their functional relationships – including positional dependence, sequence context, and higher-order interactions – that are causally-linked to model predictions. Hence, this is a powerful tool to quantitatively interpret black box models in genomics.

To generate data-driven hypotheses, first-order and second-order attribution methods can be employed to identify important *local* features. Because attribution maps can be noisy, it may be beneficial to employ CNNs that are designed to learn more interpretable representations in first layer filters (Koo & Eddy, 2019). It turns out that CNNs designed to learn interpretable filters also yield more reliable representations with attribution methods (Koo et al., 2019). Clustering attribution maps with TF-MoDISco may provide another line of evidence (Shrikumar et al., 2018).

Model interpretability is a process. There is no “magic bullet” methodology that will uncover all relevant features and their complex relationships. We recommend approaching model interpretability as a good experimentalist would approach a biological problem – with steady stream of positive and negative control experiments. Carefully designed computational experiments can test alternative hypotheses and also further support scientific claims. Global importance analysis provides the necessary framework to carry out these control experiments *in silico* in a quantitative manner.

